# C9orf72-ALS mutation drives mitophagy impairments in iNeurons

**DOI:** 10.1101/2025.02.28.640849

**Authors:** James A. K. Lee, Sarah Granger, Katie Roome, Scott P Allen, Laura Ferraiuolo, Pamela J Shaw, Heather Mortiboys

## Abstract

**Introduction:** ALS is a neurodegenerative disorder characterised by progressive upper and lower motor neuron loss. A GGGGCC hexanucleotide repeat expansion (HRE) in the C9orf72 gene is the most common mutation found in populations of European descent. Mitochondrial dysfunction has been observed in C9orf72-ALS patients and models of the disease, however reports on mitochondrial clearance via mitophagy in C9orf72-ALS are limited.

**Results:** iNeurons from C9orf72-ALS patients displayed reduced mitochondrial membrane potential and reduced mitophagy, due to reductions in autophagosome production and reduced ULK1 recruitment to mitochondria. No consistent changes to PINK1/Parkin or BNIP3 mitophagy pathways were observed.

**Conclusions:** Our data show that mitochondrial function is impaired in C9orf72-ALS patient iNeurons. An in-depth characterisation of mitophagy suggests that a deficit in autophagosome production is responsible and provides further evidence that toxic gain-of-function mechanisms in C9orf72-ALS are responsible for autophagy deficits.

## Introduction

Amyotrophic lateral sclerosis (ALS) is the most common form of motor neuron disease, characterised by progressive loss of upper and lower motor neurons. Resulting symptoms include progressively developing muscle weakness and paralysis. In cohorts of European descent, the most commonly identified genetic subtype is a hexanucleotide repeat expansion in the first intron of C9orf72 (Woollacott & Mead, 2014). Multiple pathogenic mechanisms have been identified to contribute to neuron toxicity in C9orf72-ALS, including haploinsufficiency, sequestration of RNA-binding proteins at RNA foci, and generation of dipeptide repeat proteins (DPR’s) from RAN translation of the repeat expansion (DeJesus-Hernandez et al., 2011; Renton et al., 2011) These mechanisms appear to synergise to contribute to neurodegeneration, with RNA foci and DPR production thought to be the primary drivers of neuron loss and reductions in the C9orf72 protein exacerbating the pathophysiology (Braems et al., 2020).

There is a wealth of evidence describing mitochondrial dysfunction in ALS models (reviewed recently by Lee et al., 2024). Dysfunctional mitochondria have also been observed in C9orf72-ALS models, with a majority of reports suggesting increased mitochondrial fission and disrupted mitochondrial function (Choi et al., 2019; Dafinca et al., 2016; Li et al., 2020; Lopez-Gonzalez et al., 2016; Mehta et al., 2021; Onesto et al., 2016; Petrozziello et al., 2022). We have also previously shown increases in mitochondrial reactive oxygen species (ROS) production in C9orf72-ALS iNeurons, a finding corroborated in *Drosophila* models of C9orf72-ALS (Au et al., 2024). These dysfunctional mitochondria may accumulate in cells due to impaired clearance via mitophagy. Reductions in mitophagy have been reported in *Drosophila* expressing arginine-rich DPR’s and in zebrafish models combining C9orf72 loss and polyGP expression, however these findings have not yet been confirmed in patient-derived neuronal models (Au et al., 2024; de Calbiac et al., 2024).

This study utilises neuronal progenitor cells directly reprogrammed from patient fibroblasts, allowing us to retain the genetic background and age phenotype of donor cells (Gatto et al., 2021; Meyer et al., 2014). We have previously shown that generation of iNeurons from iNPC’s results in cells expressing neuronal markers β-III tubulin, MAP2 and NeuN (Au et al., 2024). The present study includes an extensive characterisation of mitochondria and mitophagy in a patient-derived model of C9orf72-ALS, building on previous work showing disruptions to mitophagy in neurons of other ALS genotypes. We show iNeurons carrying the C9orf72-HRE mutation are deficient in mitophagy, with reductions in autophagosome generation. In contrast to other ALS genotypes, no defects in PINK1/Parkin-dependent and no consistent changes in BNIP3/BNIP3L-dependent mitophagy were observed. Modulation of ULK1 did not rescue mitophagy or autophagy deficits. This study provides evidence that disruption to the autophagy machinery produces deficits in mitophagy in C9orf72-ALS, with evidence that toxic gain-of-function mechanisms are responsible for autophagy disruption.

## Methods

### iNPC tissue culture, iNeuron differentiation and compound treatments

Induced Neuronal Progenitor Cells (iNPC’s) were generated as previously described (Meyer et al., 2014). Table 1 includes details for patient lines used in this study. The generation and use of cells derived from these iNPC’s has been published before in Meyer et al., 2014 and Au et al, 2024. iNPC’s were maintained in DMEM: F12 with GlutaMAX (Gibco), supplemented with 1% N2 (Invitrogen), 1% B27 (Invitrogen) and 20ng/ml FGF-Basic (PeproTech). Cells were grown on fibronectin (R&D Biosystems) coated cell culture dishes and routinely sub-cultured every 2-3 days using Accutase (Corning) to detach them.

**Table 1:**
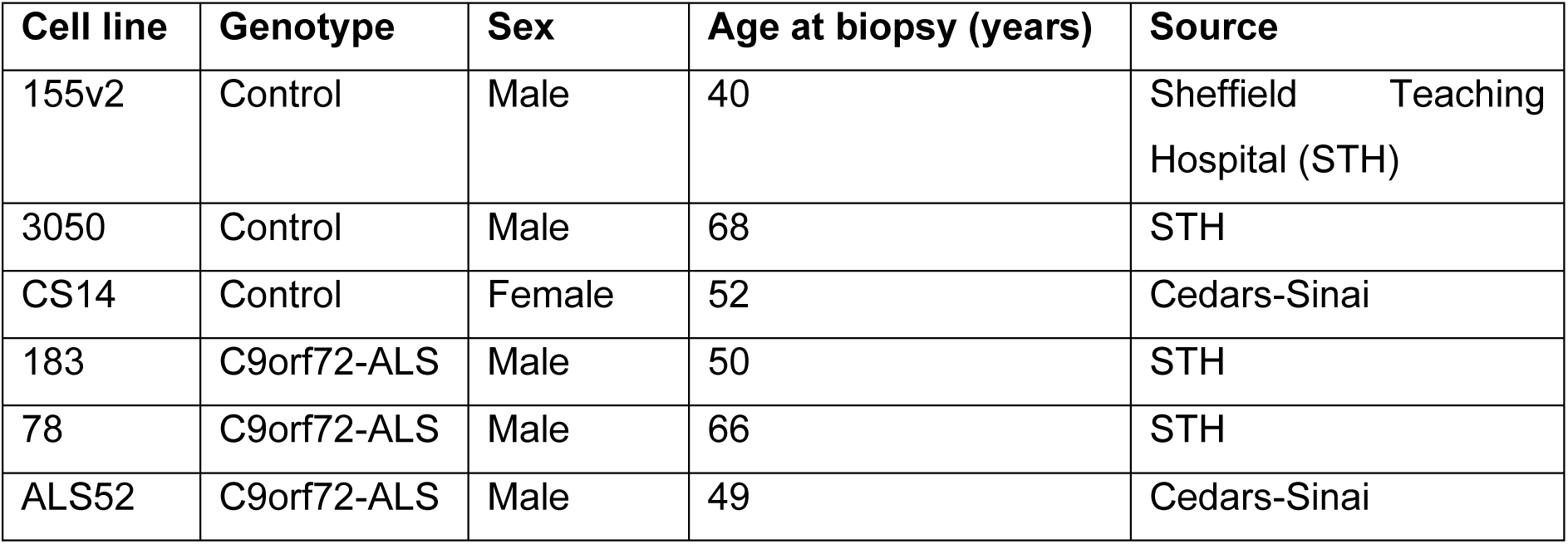
Cell lines used in this study.

| Cell line | Genotype | Sex | Age at biopsy (years) | Source |
| --- | --- | --- | --- | --- |
| 155v2 | Control | Male | 40 | Sheffield Teaching Hospital (STH) |
| 3050 | Control | Male | 68 | STH |
| CS14 | Control | Female | 52 | Cedars-Sinai |
| 183 | C9orf72-ALS | Male | 50 | STH |
| 78 | C9orf72-ALS | Male | 66 | STH |
| ALS52 | C9orf72-ALS | Male | 49 | Cedars-Sinai |

To achieve neuronal differentiation, iNPC’s were plated into 6 well plates and allowed to reach ∼80% confluency. iNeuron differentiation was started by switching to DMEM: F12 media with GlutaMAX supplemented with 1% N2, 2% B27 and 2.5μM DAPT (Sigma). After 48 hours, DAPT was removed and replaced in media with 1μM retinoic acid (Sigma), 1μM smoothened agonist (Millipore) and 2.5μM Forskolin (Cayman Chemical). Cells were maintained in this media for 16 days before being used in assays.

To assess dependency of neurons on OXPHOS or glycolysis for ATP production throughout differentiation, cells were treated with 10μM oligomycin to inhibit OXPHOS, 50mM 2-deoxyglucose to inhibit glycolysis, or both simultaneously for 30 mins at 37°C, before ATP measurements were performed with the ATPlite Luminescence Assay as per the manufacturers instructions (Revvity). From this we can calculated the percentage of ATP levels setting untreated at each time point to 100%.

For deferiprone treatments, cells were treated with either 500μM or 1mM for 24 hours prior to fixing cells and immunostaining. For oligomycin/antimycin A treatments, cells were treated with 10μM oligomycin and 4μM antimycin A for 1 hour before fixing cells and immunostaining. For nilotinib, BL-918 and A769662 treatments, cells were treated with 1μM or 5μM nilotinib/BL918, or 1μM or 10μM A769662, for 24 hours prior to fixing cells and immunostaining. For autophagy induction, cells were treated with either 200nM bafilomycin A1, 1μM rapamycin, 15μM chloroquine or 1μM rapamycin/15μM chloroquine in combination for 5 hours before fixing cells and immunostaining.

### Live imaging of iNeurons

For mitochondrial membrane potential (MMP) measurements, cells were stained for 1 hour with 20μM Hoechst, 80nM TMRM and 200nM MitoTracker Green diluted in MEM. Cells were imaged using an Opera Phenix high content imaging system (Revvity) with a 40x water immersion objective across 15 fields of view and 6 Z stacks. Cells were maintained at 37℃ and 5% CO2 throughout imaging. Image analysis was performed using a custom protocol in Harmony 4.9 (Revvity).

For live mitophagy flux measurements, cells were stained for 1 hour with 20μM Hoechst, 80nM tetramethylrhodamine methyl ester (TMRM), 50nM Lysotracker DeepRed and 200nM MitoTracker Green diluted in MEM. For induced cells, 10μM oligomycin and 4μM Antimycin A were added immediately before imaging. Cells were imaged using an Opera Phenix high content imaging system (Revvity) with a 40x water immersion objective across 9 fields of view and 4 Z stacks, with imaging repeated every 30 minutes for 90 minutes. Cells were maintained at 37℃ and 5% CO2 throughout imaging. Image analysis was performed using a custom protocol in Harmony 4.9 (Revvity).

### Respirometry measurements

iNeurons were plated at 20,000 cells per well in a fibronectin-coated 96-well Seahorse cell culture plate (Agilent) in iNeuron media and incubated overnight at 37°C and 5% CO_2_. The following day, media was replaced with Seahorse media pH 7.4 (Agilent) and cells were incubated at 37°C in a non-CO_2_ incubator for 60 minutes prior to beginning oxygen consumption readings. The effect on oxygen consumption rate (OCR) was measured 4 times for 2.5 minutes each in the absence and presence of 1.5μM oligomycin (to determine coupled respiration), 1.5μM CCCP (to determine maximal respiration and spare respiratory capacity), and finally 0.5μM rotenone in combination with 0.5μM antimycin A (to determine proton leak and non-mitochondrial oxygen consumption). Cells were fixed in 4% PFA in PBS after the assay, stained with 10μM Hoechst for 5 mins and imaged on an InCell Analyser (GE Healthcare) to determine cell numbers per well for normalisation.

### Immunofluorescent staining

On day 18 of differentiation, cells were fixed in 4% PFA for 30 minutes. After PBS washes, cells were permeabilised in PBS with 1% Tween (PBST) and 0.1% Triton X-100 for 10 minutes. Cells were blocked in 5% horse serum in PBST for 1 hour and incubated with primary antibodies in PBST overnight at 4°C. Cells were washed in PBST and incubated with secondary antibodies for 1 hour at room temperature. Nuclei were stained with 1μM Hoechst for 5 minutes prior to imaging. Cells were imaged using an Opera Phenix high content imaging system (Revvity) with a 40x water immersion objective across 20 fields of view and 6 Z stacks. Details of primary and secondary antibodies are provided in Table 2.

**Table 2:** List of antibodies used in this study.

| **Antibody** | **Species** | **Dilution used** | **Supplier** |
| --- | --- | --- | --- |
| TOM20 | Mouse | 1:1,000 | BD Biosciences (612278) |
| LC3 | Rabbit | 1:1,000 | MBL(PM026) |
| LAMP2 | Mouse | 1:1,000 | Santa Cruz (sc-18822) |
| P62 | Rabbit | 1:1,000 | Proteintech (18420-1-AP) |
| NDP52 | Rabbit | 1:1,000 | Proteintech (12229-1-AP) |
| BNIP3 | Rabbit | 1:200 (immunostaining)<br>1:1,000 (Western blot) | Abcam (ab109362) |
| NIX | Rabbit | 1:250 (immunostaining)<br>1:1,000 (Western blot) | Abcam (8399) |
| Phospho-NIX<br>(serine 81) | Rabbit | 1:1,000 | Abcam (ab208190) |
| HSP60 | Chicken | 1:5,000 | Thermofisher (PA5-143751) |
| ULK1 | Mouse | 1:200 | Santa Cruz (sc390904) |
| C9orf72 | Rabbit | 1:1,000 | Atlas (HPA023873) |
| Tubulin (alpha) | Mouse | 1:5,000 | Invitrogen (62204) |
| GAPDH | Mouse | 1:2,000 | Proteintech (60004-1) |
| Anti-mouse Alexa<br>488 secondary<br>antibody | Goat | 1:1,000 | A32723 |
| Anti-rabbit Alexa<br>568 secondary<br>antibody | Donkey | 1:1,000 | A11037 |
| Anti-chicken Alexa<br>488 secondary<br>antibody | Goat | 1:1,000 | A11039 |
| Anti-rabbit HRP<br>secondary antibody | Goat | 1:5,000 | Dako (P044801-2) |
| Anti-mouse HRP<br>secondary antibody | Goat | 1:10,000 | Abcam (ab97040) |

### Immunoblotting

Day 18 iNeuron pellets were resuspended in 50μl lysis buffer consisting of RIPA buffer, protease inhibitor cocktail (Sigma) and phosphatase inhibitor cocktail (Sigma). After 30 minutes incubating on ice, pellets were centrifuged at 13,000rpm for 20 minutes at 4 °C. Supernatant was collected and protein content was determined using a Bradford assay as per the manufacturer’s instructions. All samples were denatured at 95°C for 5 minutes in Laemmli buffer. 20μg of protein was loaded on 7.5% or 12% SDS-polyacrylamide gels for resolving, with protein electrophoresis performed using Mini-PROTEAN Tetra Handcast system (Bio-Rad). Proteins were transferred to PVDF membranes (Millipore) at 250mA for 60 minutes. Membranes were blocked in 5% milk or BSA in tris buffered saline with Tween20 (TBST). Details of primary and secondary antibodies are provided in Table 2.

## Statistical analysis

Statistical analyses were performed in Prism 10.0 (GraphPad). Normality of data was confirmed with Shapiro-Wilk test. If data were normally distributed, two-tailed unpaired t-tests or two-way ANOVAs with Tukey’s multiple comparisons test were performed. A p-value of less than 0.05 was considered significant. If data were non-normally distributed, Mann-Whitney U tests were performed.

## Results

### C9orf72-HRE reduces mitochondrial function

We first sought to understand mitochondrial function in C9orf72-ALS iNeurons. We have previously shown that this iNeuron differentiation protocol yields neurons that stain positive for pan-neuronal markers β-III tubulin, MAP2 and NeuN, resulting in 83% of neurons at day18 staining positive for β-III tubulin, 75% positive for MAP2 and 52% positive for NeuN (Au et al., 2024), indeed the neurons used in Au et al, 2024 were generated at the same time from the same lines as those used in this study. To assess the relative contribution of glycolysis and OXPHOS to ATP levels in the iNeurons, we measured ATP levels after treatment with inhibitors of glycolysis (2-deoxyglucose) and OXPHOS (oligomycin). In agreement with previous reports, iNPC’s were found to be heavily glycolytic, with mitochondrial inhibition only reducing ATP levels by ∼19% relative to vehicle treated controls (Schwartzentruber et al., 2020). By the end of differentiation, oligomycin led to larger reductions in ATP levels, down to 55% of UT levels in control iNeurons and 61% in C9orf72 iNeurons (Figure 1A). Therefore, these iNeurons have begun the metabolic switch, however they are still reliant on glycolysis for 50% of their ATP production. No significant difference was observed in ATP levels between control and C9orf72-ALS iNeurons (not shown).

**Figure 1:**
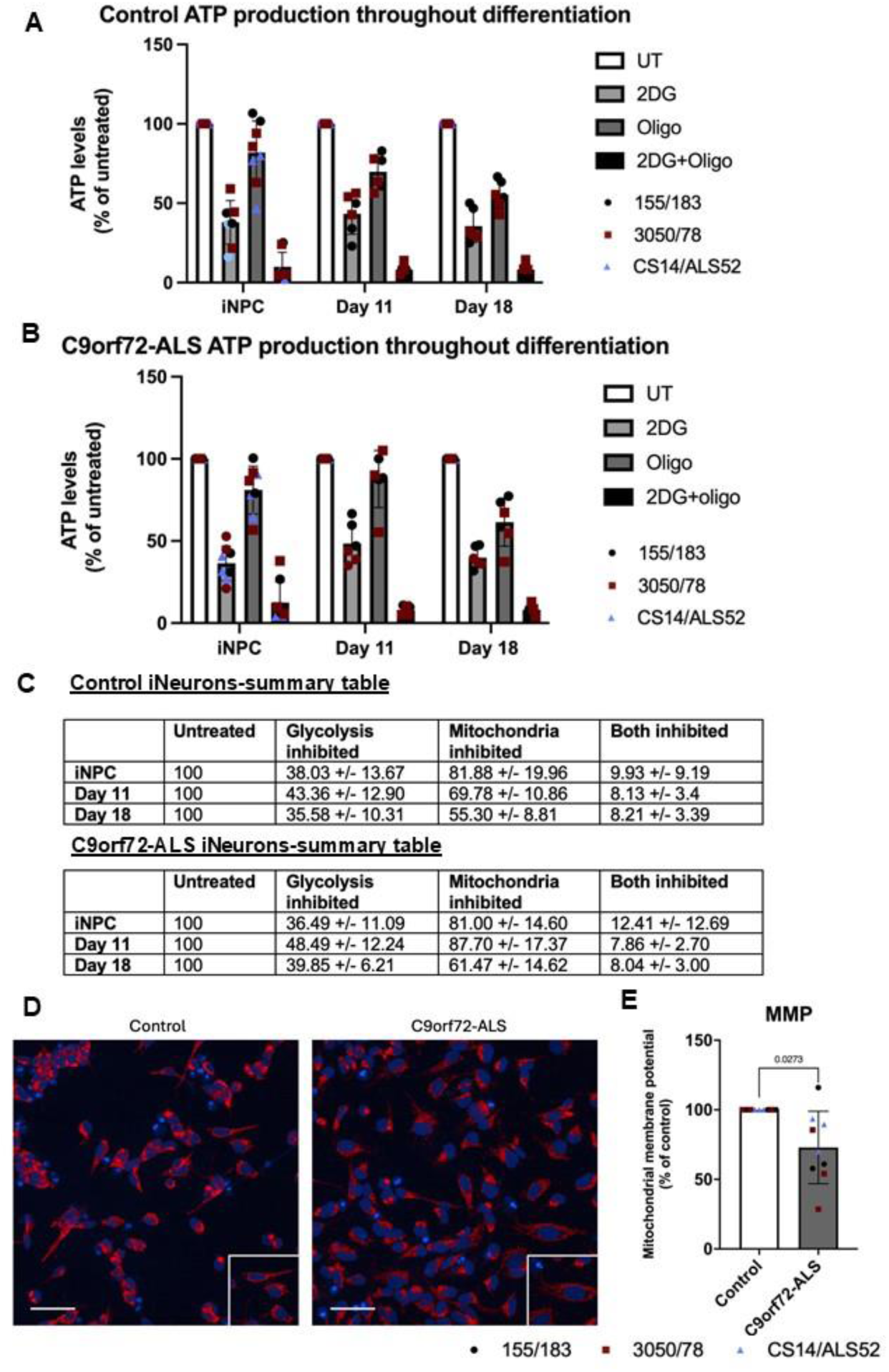
Deficits in mitochondrial function in C9orf72-ALS iNeurons. **(A-B)** ATP production dependent on glycolysis and oxidative phosphorylation throughout differentiation in control (A) and C9orf72-ALS (B) iNeurons. **(C)** Summary tables indicating percentage of ATP produced after inhibition of glycolysis or oxidative phosphorylation. **(D)** Representative images of control and C9orf72-ALS iNeurons stained with mitochondrial membrane potential marker TMRM. Scale bar= 100μM. **(E)** Quantification of mitochondrial membrane potential (mean ±SD, Wilcoxon test). All quantification was performed on 3 different differentiations of 2-3 control and C9orf72-ALS iNeuron lines, each data point represents one unique differentiation of a control/C9orf72-ALS line, taken from a mean of approximately 100-500 cells. Data from each patient/control pair are uniquely colour-coded.

Next, we stained iNeurons with TMRM and measured fluorescence intensity as an indicator of mitochondrial membrane potential (MMP). We observed a significant decrease in MMP (Figure 1B-C). We simultaneously measured mitochondrial morphology using a membrane potential independent dye and using mitochondrial staining of TOM20, however no differences in area, length, width: length or roundness were observed (Figure S1). To further interrogate mitochondrial function, we used the Seahorse Bioanalyser to investigate mitochondrial oxygen consumption under basal and stressed conditions. Despite variability between patient lines, we observed no consistent differences in basal OCR, proton leak, coupled respiration, maximal respiration or spare respiratory capacity (Figure S2). Given the iNeurons used in this study still relied on glycolysis for over 50% of ATP production, we also assessed extracellular acidification rate (ECAR) alongside OCR during this assay to provide an indication of glycolytic rate in these iNeurons. We found no significant differences in basal glycolytic rate, glycolytic capacity or glycolytic reserve in C9orf72-ALS iNeurons relative to controls (Figure S2).

### C9orf72-HRE leads to reduced mitophagy

Initial assessments of mitophagy were performed by staining for the mitochondrial marker TOM20 and the autophagosomal marker LC3 in fixed iNeurons. A significant decrease in mitochondria co-localising with autophagosomes was observed in C9orf72-ALS iNeurons relative to controls (Figures 2A and B). In contrast, when we investigated mitochondria-lysosome co-localisation in iNeurons using live imaging, under basal conditions and after induction with oligomycin-antimycin A, we observed no differences in the number of mitochondria colocalising with lysosomes (Figure 2 C-E).

**Figure 2:**
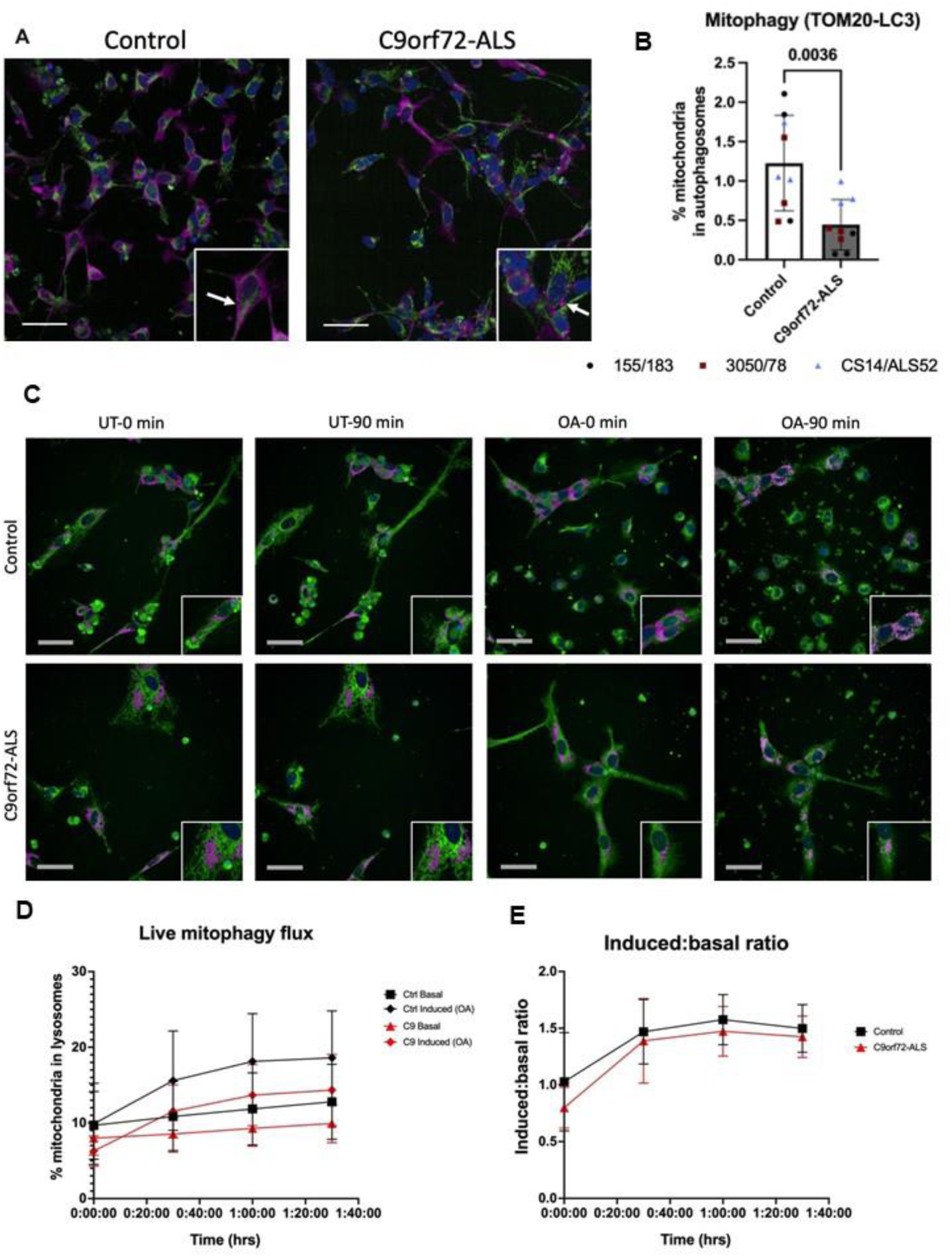
Mitophagy deficit in C9orf72-ALS iNeurons. **(A)** Representative images of control and C9orf72-ALS iNeurons stained with mitochondrial marker TOM20 (green) and autophagosomal marker LC3 (magenta). Scale bar= 100μM. Arrows indicate examples of TOM20-LC3 co-localisation. **(B)** Quantification of percentage of mitochondria co-localising with autophagosomes (mean ±SD, unpaired t-test). Each data point represents one unique differentiation of a control/C9orf72-ALS line, taken from a mean of approximately 100-500 cells. **(C)** Representative images of control and C9orf72-ALS iNeurons during mitophagy flux assay in untreated (UT) and induced (OA) conditions. Mitochondria in green, lysosomes in magenta. Scale bar= 100μM. **(D-E)** Quantification of live mitophagy flux assay and induced: basal ratio. All quantification was performed on 3 different differentiations of 2-3 control and C9orf72-ALS iNeuron lines. Data from each patient/control pair are uniquely colour-coded.

### C9orf72-HRE disrupts autophagy machinery

There have been multiple reports indicating a role for the C9orf72 protein and DPR’s on autophagy initiation and autophagosome-lysosome fusion. We observed no differences in autophagosome-lysosome co-localisation in iNeurons, however we did observe a reduction in autophagosome density in C9orf72-ALS iNeurons (Figure 3C). We sought to confirm whether autophagy induction was impaired in these iNeurons by manipulating autophagy with well characterised autophagy modulators (Klionsky et al., 2021). iNeurons were treated with bafilomycin A1 or chloroquine, both of which block lysosome degradation of autophagosomes and their contents. Rapamycin was also used as an inhibitor of mTOR, to induce autophagy and autophagosome production. Treatments with bafilomycin, rapamycin and chloroquine led to smaller increases in autophagosomes in C9orf72 iNeurons relative to controls (Figure 3D).

**Figure 3:**
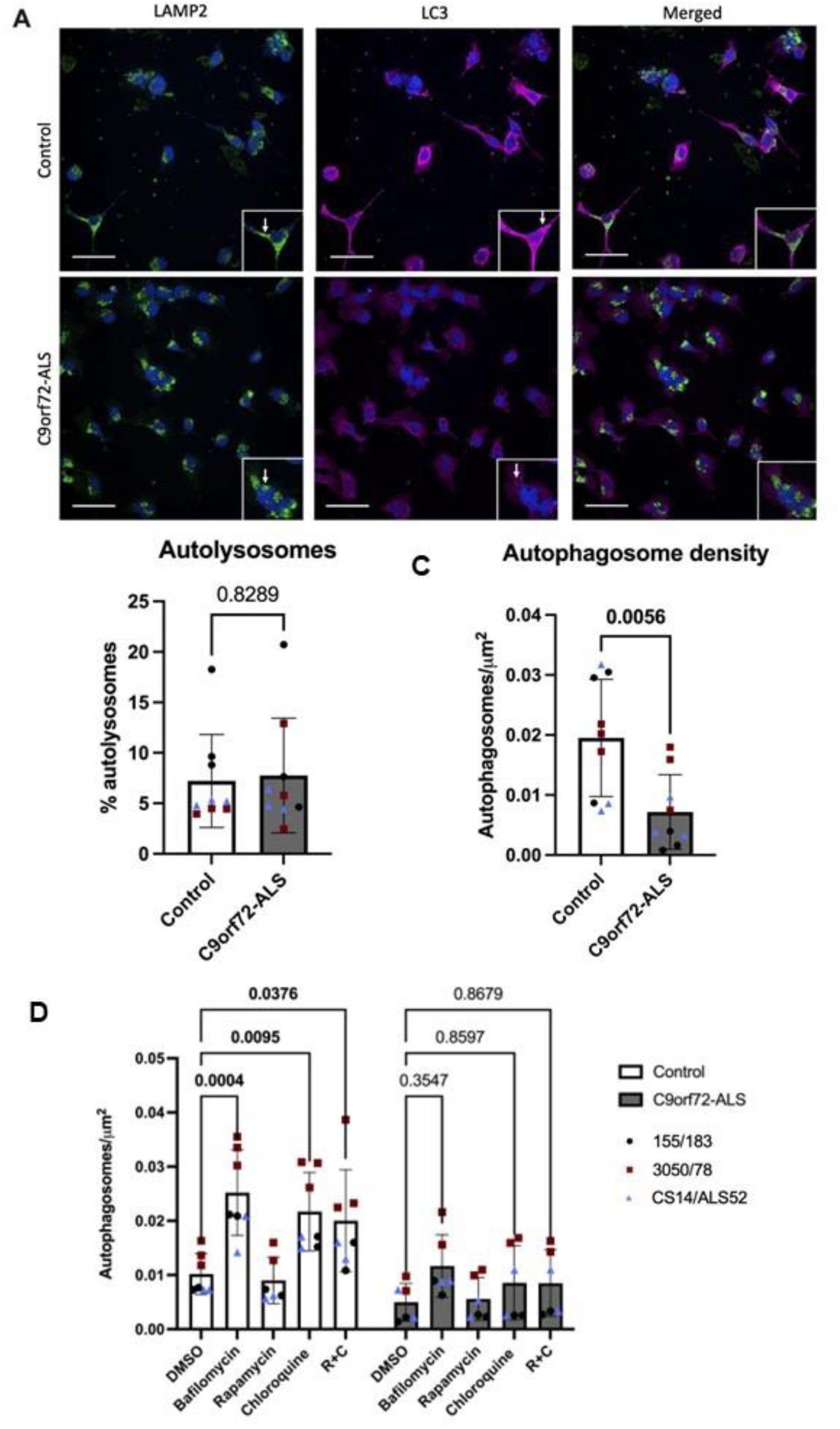
Autophagosome deficit in C9orf72-ALS iNeurons. **(A)** Representative images of control and C9orf72-ALS iNeurons stained with lysosomal marker LAMP2 (green) and autophagosomal marker LC3 (magenta). Arrows indicated example lysosomes/autophagosomes on respective images. Scale bar= 100μM. **(B)** Quantification of percentage of autophagosomes co-localising with lysosomes (mean ±SD, unpaired t-test). **(C)** Quantification of autophagosome density (mean ±SD, unpaired t-test). **(D)** Quantification of autophagosome density after treatment with bafilomycin, rapamycin and chloroquine (mean ±SD, 2-way ANOVA with Tukey’s multiple comparisons test). Each data point represents one unique differentiation of a control/C9orf72-ALS line, taken from a mean of approximately 100-500 cells. All quantification was performed on 3 different differentiations of 3 control and C9orf72-ALS iNeuron lines. Data from each patient/control pair are uniquely colour-coded.

### PINK1-Parkin and BNIP3/BNIP3L mitophagy pathways are not consistently dysregulated in C9orf72-ALS

We next sought to establish whether common mitophagy initiating pathways upstream of autophagosomal engulfment were disrupted in C9orf72-ALS. Previous studies have shown disruptions to PINK1/Parkin, BNIP3 and BNIP3L-dependent mitophagy in SOD1-ALS, TDP43-ALS, FUS-ALS and sporadic ALS (sALS) neurons (Lagier-Tourenne et al., 2012; Palomo et al., 2018; Rogers et al., 2017; Stribl et al., 2014). We observed no differences in Parkin co-localisation with mitochondria in C9orf72-ALS iNeurons relative to controls, although there was large variation between the *C9orf72*-ALS patient lines. Downstream of Parkin, we also observed no differences in phospho-ubiquitin accumulation at mitochondria in C9orf72-ALS iNeurons, although there was large variation between the *C9orf72*-ALS patient lines (Figure 4). The recruitment of Sequestosome-Like Receptor proteins NDP52 and P62 were also not disrupted in C9orf72-ALS iNeurons (Figure S5). Co-localisation of BNIP3 and BNIP3L with mitochondria basally showed a trend towards being reduced however due to large variability between both controls and patients this was not significant (Figure 5). After induction with the iron chelator deferiprone (DFP), BNIP3 and BNIP3L localisation at the mitochondria was unaffected in C9orf72-ALS iNeurons (Figure 5). Moreover, no changes in expression of BNIP3 and BNIP3L were observed via immunoblot (Figure S4). Interestingly, we also observed a significant negative correlation between mitochondrial co-localisation and protein expression levels in BNIP3L but not BNIP3 in our iNeuron lines (Figure S4 G-H).

**Figure 4:**
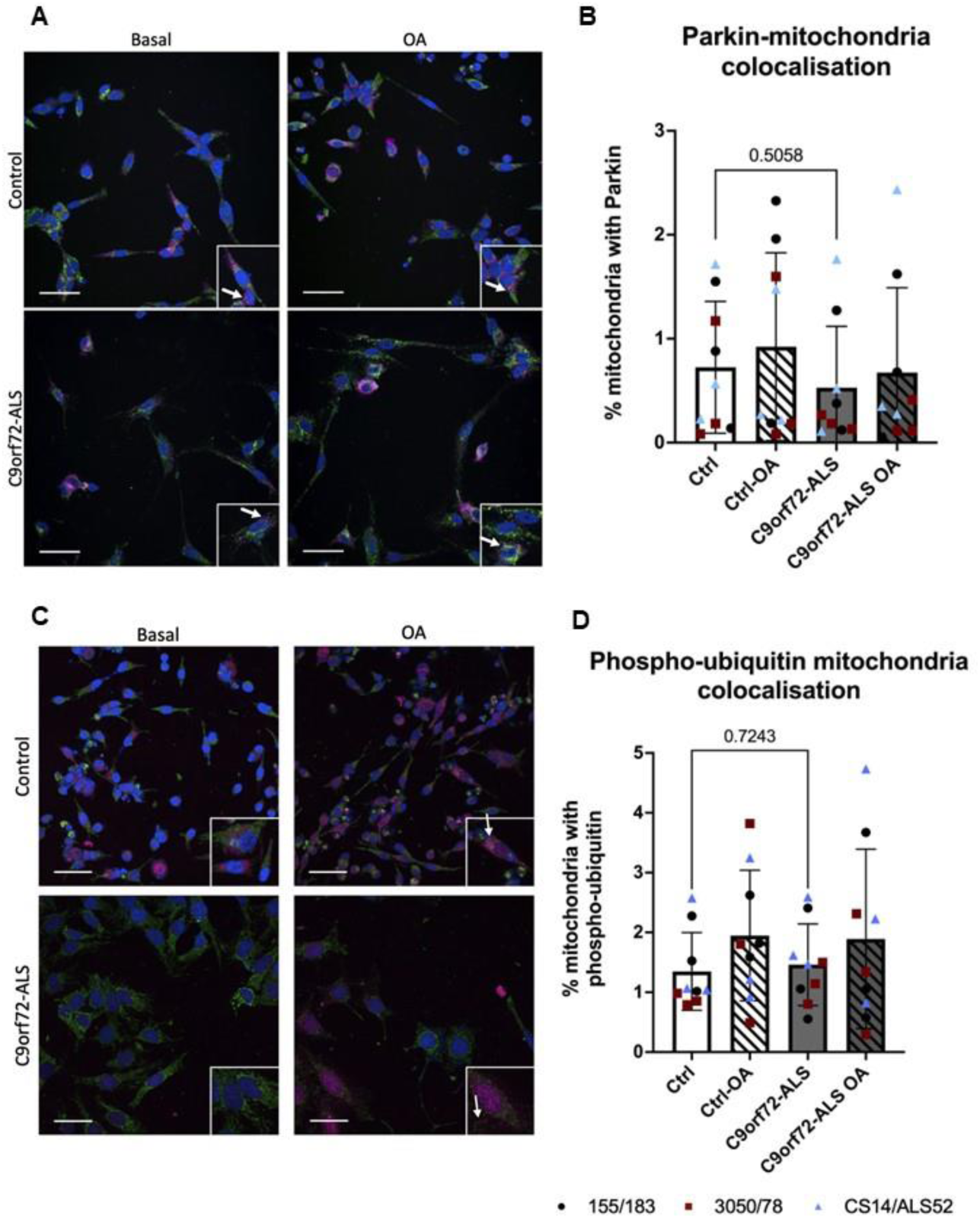
Parkin-dependent mitophagy is unaffected in C9orf72-ALS iNeurons. **(A)** Representative images of control and C9orf72-ALS iNeurons stained with mitochondrial marker TOM20 (green) and Parkin (magenta), under basal conditions and after mitophagy induction with oligomycin/antimycin A (OA) treatment. Arrows indicated example mitochondria staining with Parkin. Scale bar= 100μM. **(B)** Quantification of percentage of mitochondria staining with Parkin (mean ±SD, unpaired t-test). Each data point represents one unique differentiation of a control/C9orf72-ALS line, taken from a mean of approximately 100-500 cells. **(C)** Representative images of control and C9orf72-ALS iNeurons stained with mitochondrial marker TOM20 (green) and phospho-ubiquitin (magenta), under basal conditions and after mitophagy induction with oligomycin/antimycin A (OA) treatment. Arrows indicated example mitochondria staining with phospho-ubiquitin. Scale bar= 100μM. **(D)** Quantification of percentage of mitochondria staining with phospho-ubiquitin (mean ±SD, unpaired t-test). Each data point represents one unique differentiation of a control/C9orf72-ALS line, taken from a mean of approximately 100-500 cells. All quantification was performed on 3 different differentiations of 3 control and C9orf72-ALS iNeuron lines. Data from each patient/control pair are uniquely colour-coded.

**Figure 5:**
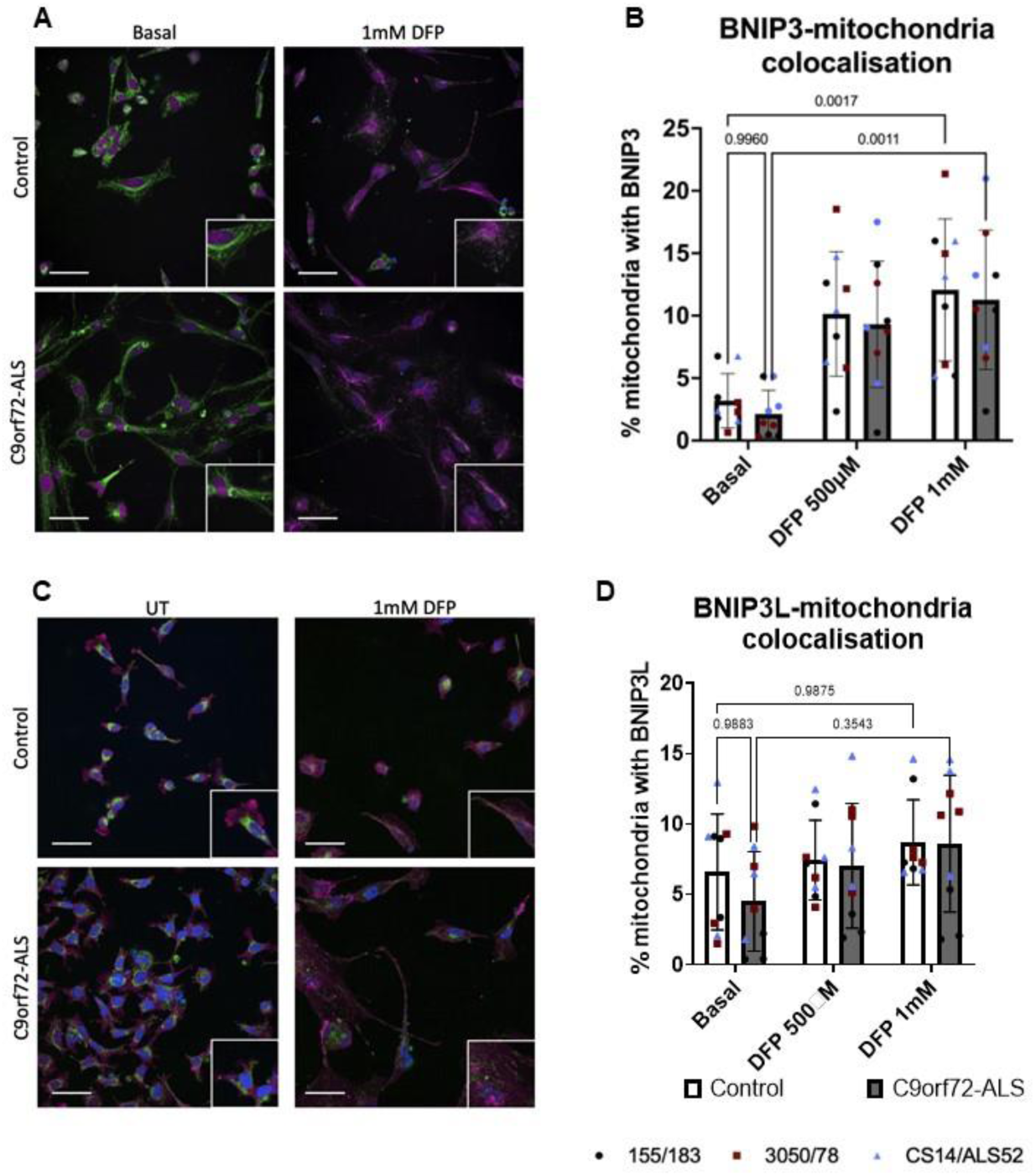
BNIP3-dependent and BNIP3L-dependent mitophagy is unaffected in C9orf72-ALS iNeurons. **(A)** Representative images of control and C9orf72-ALS iNeurons stained with mitochondrial marker TOM20 (green) and BNIP3 (magenta), under basal conditions and after mitophagy induction with deferiprone (DFP) treatment. Scale bar= 100μM. **(B)** Quantification of percentage of mitochondria staining with BNIP3 (mean ±SD, two-way ANOVA). Each data point represents one unique differentiation of a control/C9orf72-ALS line, taken from a mean of approximately 100-500 cells. **(C)** Representative images of control and C9orf72-ALS iNeurons stained with mitochondrial marker TOM20 (green) and NIX (magenta), under basal conditions and after mitophagy induction deferiprone (DFP) treatment. Scale bar= 100μM. **(D)** Quantification of percentage of mitochondria staining with NIX (mean ±SD, two-way ANOVA). Each data point represents one unique differentiation of a control/C9orf72-ALS line, taken from a mean of approximately 100-500 cells. All quantification was performed on 2-3 different differentiations of 3 control and C9orf72-ALS iNeuron lines. Data from each patient/control pair are uniquely colour-coded.

### ULK1 recruitment is disrupted in C9orf72-ALS iNeurons

C9orf72 has previously been shown to mediate the recruitment of the ULK1 complex to the early autophagosome (Sellier et al., 2016; Webster et al., 2016). We investigated co-localisation of mitochondria with ULK1 to investigate whether its recruitment to mitochondria prior to mitophagy was disrupted. We observed a significant reduction in ULK1 puncta co-localising with mitochondria (Figure 6). We also investigated whether increasing ULK1 activity would rescue autophagy deficits caused by this impaired recruitment. Treatment with the AMPK activator A769662 and nilotinib, a drug that has been previously shown to activate ULK1, had no impact on mitophagy or autophagosome production in control of C9orf72 iNeurons. However, the ULK1 activator BL-918 was able to increase mitophagy in control at the highest concentration but not in C9orf72-ALS iNeurons (Figure S7). Moreover, we show no significant changes in C9orf72 expression in C9orf72-ALS iNeurons (Figure S4).

**Figure 6:**
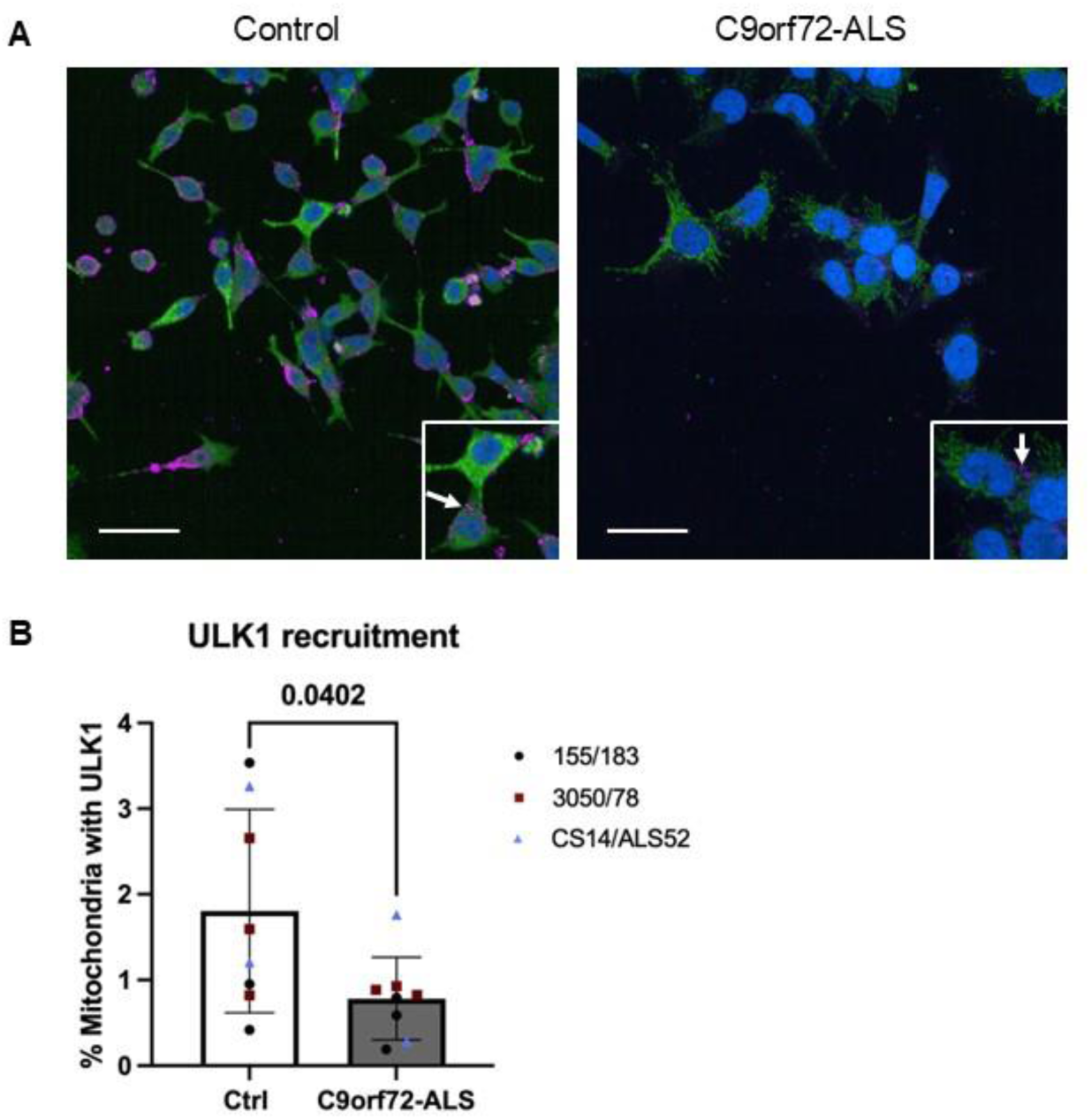
ULK1 recruitment to mitochondria is reduced in C9orf72-ALS iNeurons. **(A)** Representative images of control and C9orf72-ALS iNeurons stained with mitochondrial marker HSP60 (green) and ULK1 (magenta). Scale bar= 100μM. Arrows indicated example mitochondria staining with ULK1. **(B)** Quantification of percentage of mitochondria staining with ULK1 (mean ±SD, unpaired t-test). Each data point represents one unique differentiation of a control/C9orf72-ALS line, taken from a mean of approximately 100-500 cells. All quantification was performed on 2-3 different differentiations of 3 control and C9orf72-ALS iNeuron lines. Data from each patient/control pair are uniquely colour-coded.

## Discussion

Multiple reports have indicated disrupted autophagy in models of C9orf72-ALS (Beckers et al., 2023; Sellier et al., 2016; Webster et al., 2016). Despite this, the potential impact of C9orf72-HRE mutations on mitophagy remains poorly understood. To address this, we have performed for the first time a comprehensive characterisation of mitophagy in C9orf72-ALS iNeurons. Our findings indicate no disruption to mitophagy initiating pathways and suggest that disruption of the autophagy machinery is responsible for reduced mitophagy observed in C9orf72-ALS.

Prior to this study, the impact of C9orf72-ALS on mitophagy was poorly understood. Ectopic expression of the G4C2-HRE in *Drosophila* has previously been shown to reduce mitophagy (Au et al., 2024). However, the cellular mechanisms driving this mitophagy deficit were not explored. There have been multiple reports indicating disruptions to PINK1/Parkin-mitophagy and BNIP3/BNIP3L-dependent mitophagy in TDP43-ALS, SOD1-ALS and sALS in iPSC-neurons, mouse neurons and patient peripheral blood mononuclear cells (PBMC’s) (Araujo et al., 2020; Lagier-Tourenne et al., 2012; Palomo et al., 2018; Rogers et al., 2017; Stribl et al., 2014). Equally, despite conflicting reports, the contribution of the C9orf72-HRE to disrupting autophagy machinery has also been previously described, with reports indicating changes to lysosome function, autophagosome-lysosome fusion and autophagosome production (Beckers et al., 2023; Sellier et al., 2016; Webster et al., 2016; Yang et al., 2016). These previous findings highlight the importance of identifying which specific areas of the mitophagic pathway might be affected to best inform attempts at therapeutic intervention.

PINK1/Parkin-dependent mitophagy is the most well characterised mitophagy pathway, typically triggered when PINK1 stabilises in the mitochondrial membrane on mitochondrial injury or depolarisation (Jin & Youle, 2012). Subsequently Parkin is recruited, leading to the accumulation of phospho-ubiquitin chains on outer mitochondrial membrane proteins. We have previously shown increases in mitochondrial ROS production in C9orf72-ALS iNeurons, and here demonstrate reduced MMP, both indicating dysfunctional mitochondria (Figure 1; Au et al., 2024). Despite this, no changes to Parkin or phospho-ubiquitin co-localisation with mitochondria were observed under basal conditions (Figure 4). Induction of mitophagy via treatment with oligomycin/antimycin A, which typically induces PINK1/Parkin-dependent mitophagy, led to similar increases in mitophagy rate in C9orf72-ALS iNeurones relative to controls (Figure 2). Previous reports have suggested PINK1/Parkin-dependent mitophagy may be disrupted in ALS, with TDP43-ALS, FUS-ALS and sALS patient derived neurons displaying reduced Parkin expression (Lagier-Tourenne et al., 2012). SOD1 and TDP43 mouse models also display reductions in Parkin expression (Stribl et al., 2014; Rogers et al., 2017). Moreover, Parkin recruitment following mitochondrial depolarisation has previously been shown to be reduced in highly oxidative cells (Van Laar et al., 2011). Higher mitophagy rates have also been demonstrated in glycolytic cells compared to oxidative cells (Montava-Garriga et al., 2020). Furthermore, in some neuronal models, deficiencies in Parkin-dependent mitophagy have only been observed in oxidative neurons compared to glycolytic neurons (Schwartzentruber et al., 2020). Given the neurons utilised in this study still produced a significant amount of their ATP through glycolysis, this may have influenced observed phenotypes. These findings warrant further investigation in patient-derived models to determine to what extent Parkin-dependent mitophagy may be impacted in non-C9orf72-ALS. Further experiments to determine the impact of oxidative status of the neurons on Parkin-dependent mitophagy would also be useful in these models to determine the impact of this mitophagy pathway.

BNIP3 and BNIP3L have both been identified as mitophagy receptors that act independently of ubiquitination and have been shown to regulate basal mitophagy (Elcocks et al., 2023). Previous studies have identified hypoxia and iron chelation as inducers of BNIP3 and NIX-dependent mitophagy (Allen et al., 2013; Ganley & Simonsen, 2022). We show that under basal conditions, and after induction with deferiprone, there are no differences in BNIP3 or BNIP3L recruitment to mitochondria (Figure 5). Moreover, no differences in expression of either protein were observed under basal conditions (Figure S4). However, due to variability in C9orf72-ALS iNeuron lines, further investigations in a larger cohort of patient lines would be useful to accurately establish whether the trending decrease in BNIP3L localisation to mitochondria is limited to a small number of patients or widespread in C9orf72-ALS. Interestingly, we observed a negative correlation between mitochondrial co-localisation and expression levels of BNIP3L but not BNIP3 (Figure S4 G-H). Given BNIP3L can interact with other autophagy pathways, this suggests higher levels of BNIP3L under basal conditions may be due to its interactions with other autophagy pathways, something that could warrant further investigation in these iNeurons (Li et al., 2021). We also investigated localisation of BNIP3L phosphorylated at serine-81 to mitochondria (Figure S6). Although we found no significant difference between control and C9orf72-ALS iNeurons, there were notable reductions in some C9orf72-ALS iNeuron lines relative to their matched controls (Figure S6 B). Phosphorylation of BNIP3L has been shown to enhance its function in mitophagy (Poole et al., 2021). Again, expanding the cohort size and a greater investigation of multiple phosphorylation sites could help identify if reductions in BNIP3L activity, but not expression or mitochondrial co-localisation, might impact basal mitophagy in C9orf72-ALS. The drug used to induce BNIP3/NIX-dependent mitophagy in this study, deferiprone, has been assessed in a phase I clinical trial in ALS patients with a phase II/III trial ongoing (Moreau et al., 2018). While the treatment conditions used in this study are a chronic, high dose treatment, we demonstrate increases in autophagy and mitophagy, in agreement with previous work, which may be beneficial in ALS patients if they can be achieved with lower doses of deferiprone (Allen et al., 2013).

We identified a deficit in ULK1 recruitment to mitochondria in C9orf72-ALS iNeurons and tried to circumvent this by increasing ULK1 activation by treatment with the compounds nilotinib, BL-918 and A769662. Nilotinib and A769662 have both been shown to activate AMPK leading to ULK1 activation, whilst BL-918 has been shown to activate ULK1 directly (Goransson et al., 2007; Ouyang et al., 2018; Yu et al., 2013). However, of these compounds only BL-918 could increase mitophagy and only in control iNeurons (Figure S7). This may be due to issues of target engagement, which would be needed to further understand the reason for the lack of impact of these compounds in these iNeuron models. Moreover, AMPK activation has recently been shown to enhance clearance of damaged and dysfunctional mitochondria, via Parkin-dependent mitophagy and independently of ULK1, while simultaneously reducing NIX-dependent clearance of functional mitochondria (Longo et al., 2024). These findings warrant a more detailed investigation of AMPK activation as a therapeutic target in C9orf72-ALS iNeuron models.

The morphology and quantity of lysosomes was also measured; however no significant differences were observed (Figure S3). Increases in lysosome size and number have previously been identified in C9orf72-ALS neurons and C9orf72-KO HEK293T cells (Amick et al., 2016; Beckers et al., 2023). Previous reports in C9orf72-ALS neurons do not appear to be driven by C9orf72 expression and may be caused by different levels of DPR’s between their model and the iNeuron model used in this study. Accumulation of autolysosomes has also been reported in *Drosophila* models of C9orf72-ALS expressing DPR’s (Xu et al., 2023). Although no changes in lysosome morphology were observed, changes to lysosomal function were not investigated in this study and cannot be excluded as a contributor to autophagy and mitophagy deficits.

We have identified a deficit in autophagosome production in C9orf72-ALS iNeurons, in agreement with previous reports (Webster et al., 2016; Sellier et al., 2016). Although we hypothesised that reductions in C9orf72 protein expression may be responsible, as a result of the C9orf72-HRE, we did not observe reductions in C9orf72 expression (Figure S4). These findings suggest toxic gain-of-function mechanisms associated with the C9orf72-HRE are primarily responsible for driving autophagosomal deficits and may also impact ULK1 recruitment alongside the C9orf72-SMCR8 complex as previously described (Webster et al., 2016; Sellier et al., 2016). Recent studies have indicated that toxic gain-of-function aspects of the C9orf72-HRE, such as DPR production, can contribute to autophagosome and lysosomal dysfunction in C9orf72-ALS. Autophagy deficits have been reported in patient motor neurons but not motor neurons with C9orf72 knockout (Beckers et al., 2023). Poly-PR and poly-GR production can disrupt autophagosome production by enhancing the BCL2-Beclin1 interaction, thereby inhibiting formation of the PIK3C3 complex, downstream of ULK1 in autophagy initiation (Xu et al., 2024). DPR expression in *Drosophila* neurons leads to accumulation of autolysosomes, suggesting DPR’s interfere with lysosomal degradation (Xu et al., 2023). Taken together, these findings indicate C9orf72 loss and gain of function mechanisms work separately to impair autophagy and highlight the importance of further work in models collectively studying C9orf72 mechanisms, ideally in patient-derived models, to better elucidate any potential corrective therapeutic approaches.

## Declarations

### Ethics approval

For the iNeuron work, informed consent was obtained from all participants before collection of fibroblast biopsies (Sheffield Teaching Hospital (STH) study number: STH16573, Research Ethics Committee reference: 12/YH/0330); fibroblasts obtained by Cedars-Sinai are covered by an MTA.

## Figure Legends

**Supplementary Figure 1:**
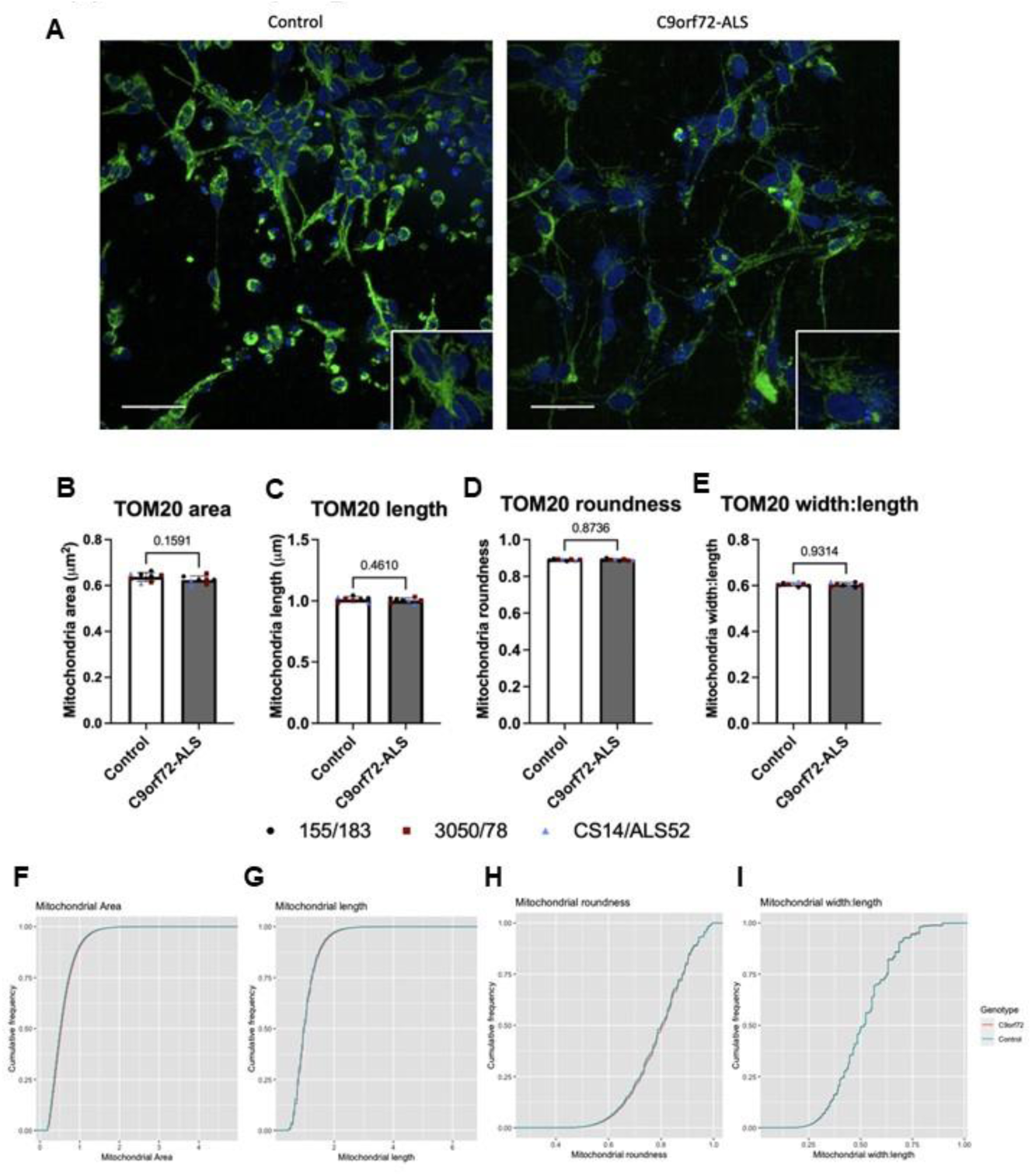
Mitochondrial morphology is unaffected in C9orf72-ALS iNeurons. **(A)** Representative images of control and C9orf72-ALS iNeurons stained with mitochondrial marker TOM20 (green). Scale bar= 100μM. **(B-E)** Quantification of segmented mitochondria area (B), length (C), roundness (D) and width:length (E) (mean ±SD, unpaired t-test). Each data point represents one unique differentiation of a control/C9orf72-ALS line, taken from a mean of approximately 100-500 cells. (F-I) Cumulative distribution functions for mitochondrial area (F), length (G), roundness (H) and width:length (I). Distributions are for each individual imaged mitochondria from approximately 100-500 cells of each unique differentiation of a control/C9orf72-ALS line, giving a total population of ∼5 million control and ∼5 million C9orf72-ALS mitochondria. All quantification was performed on 3 different differentiations of 3 control and C9orf72-ALS iNeuron lines. Data from each patient/control pair are uniquely colour-coded.

**Supplementary Figure 2:**
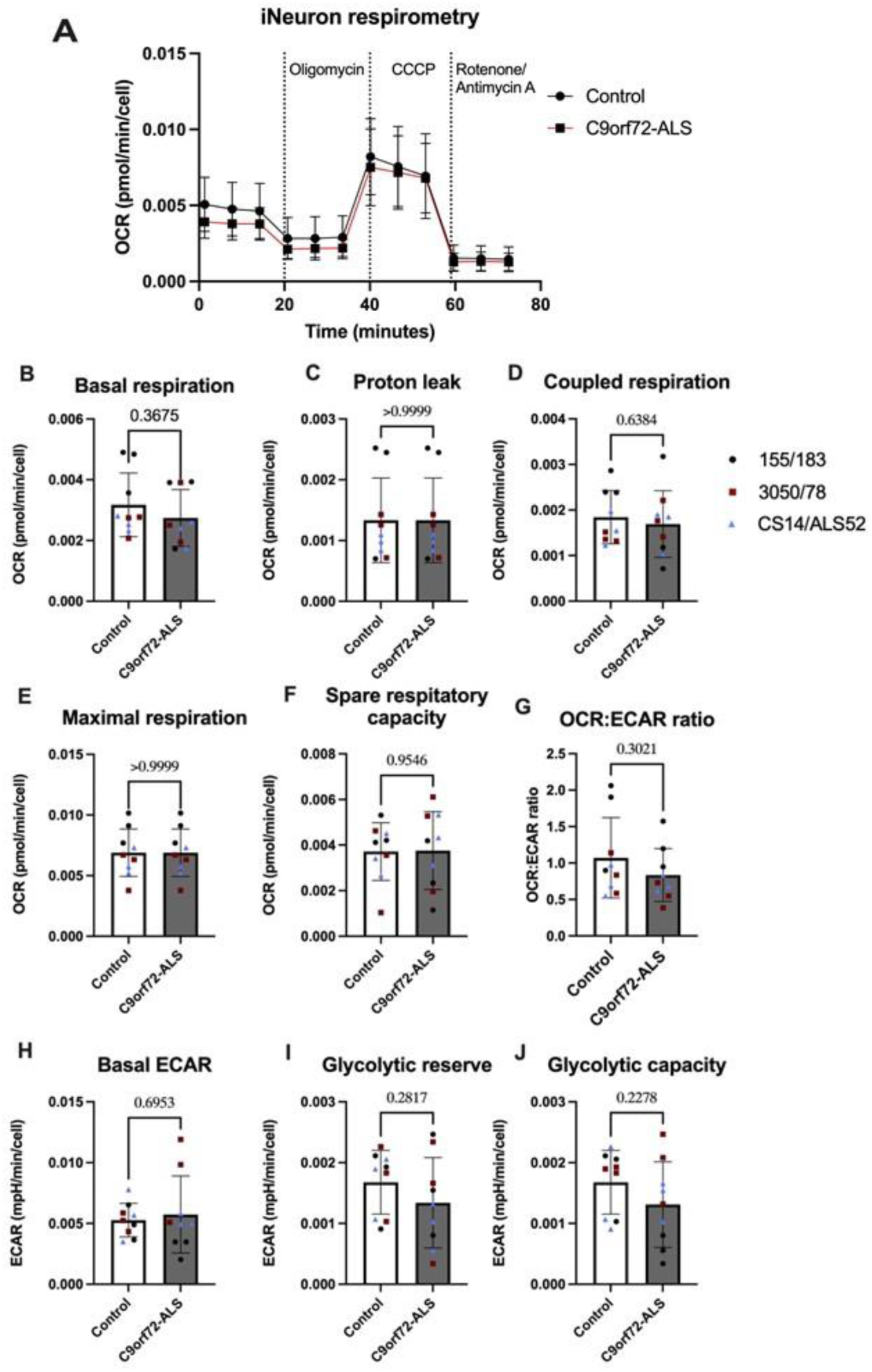
C9orf72-ALS iNeurons show no deficits in oxygen consumption. **(A)** Respirometry trace for control (black line) and C9orf72-ALS iNeurons (red line). **(B-G)** Basal respiration (B), proton leak (C), Coupled respiration (D), Maximal respiration (E), Spare respiratory capacity (F) and OCR:ECAR ratio (G) in control and C9orf72-ALS iNeurons (mean ±SD, unpaired t-test). Each data point represents one unique differentiation of a control/C9orf72-ALS line. All quantification was performed on 3 different differentiations of 3 control and C9orf72-ALS iNeuron lines. Data from each patient/control pair are uniquely colour-coded.

**Supplementary Figure 3:**
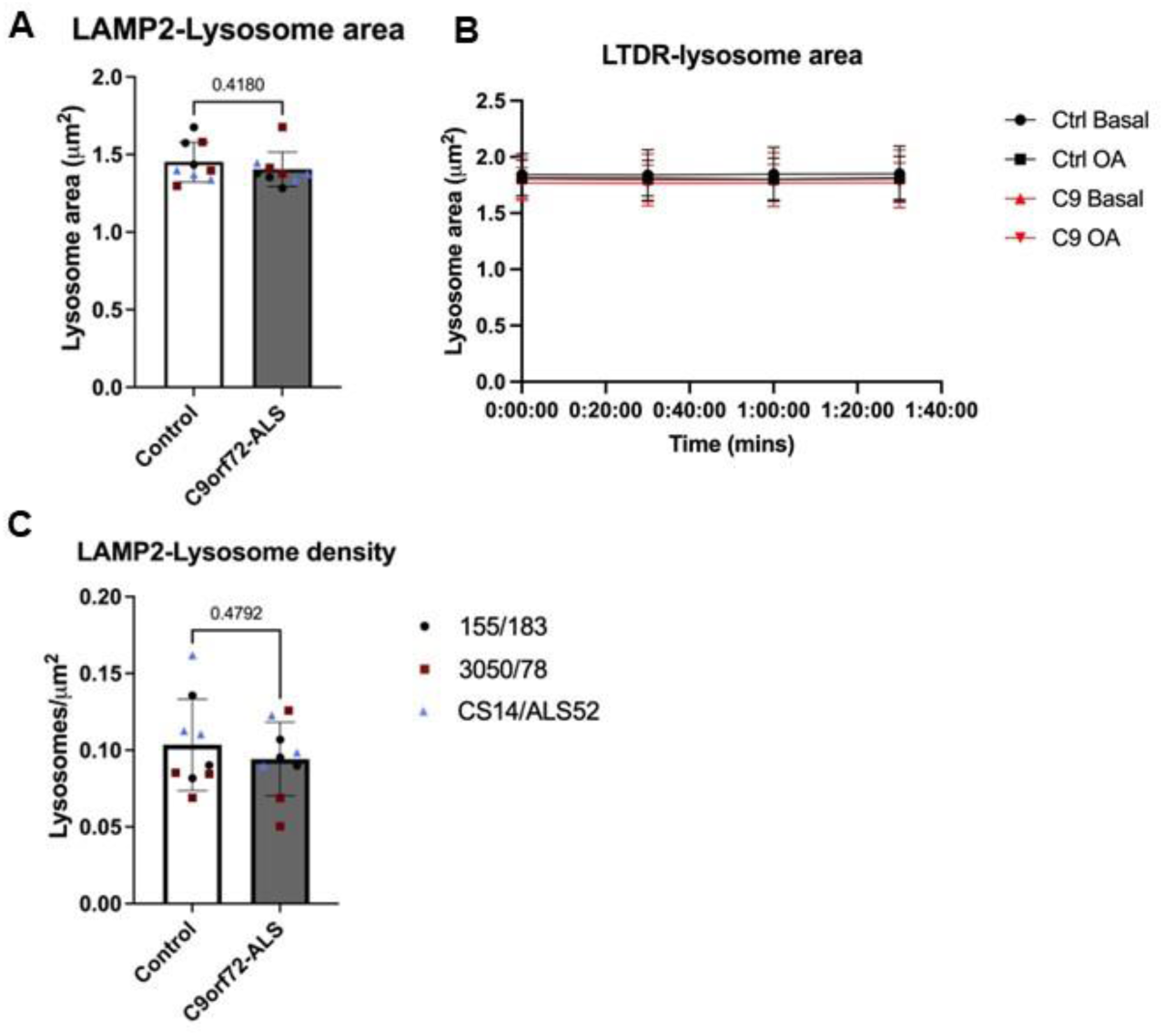
Lysosome size and density is unaffected in C9orf72-ALS iNeurons. (**A**) Quantification of lysosome area from LAMP2 staining in control and C9orf72-ALS iNeurons (mean ±SD, unpaired t-test). (**B**) Quantification of lysosome area from LysoTracker DeepRed staining during multiple timepoints of mitophagy flux assay in control and C9orf72-ALS iNeurons (mean ±SD). (C) Quantification of lysosome density from LAMP2 staining in control and C9orf72-ALS iNeurons (mean ±SD, unpaired t-test). Each data point represents one unique differentiation of a control/C9orf72-ALS line, taken from a mean of approximately 100-500 cells. All quantification was performed on 3 different differentiations of 3 control and C9orf72-ALS iNeuron lines. Data from each patient/control pair are uniquely colour-coded.

**Supplementary Figure 4:**
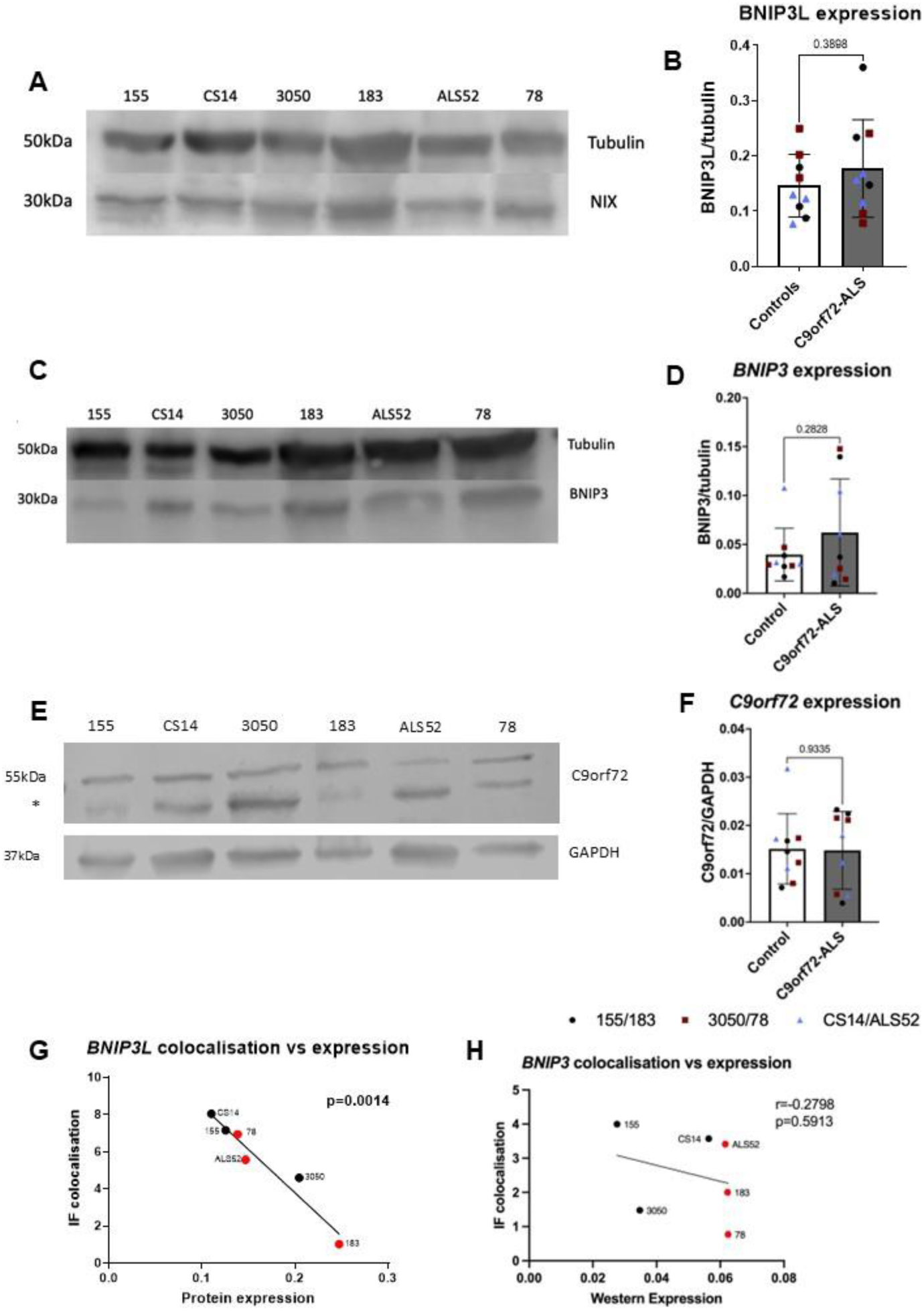
BNIP3, BNIP3L and C9orf72 expression are unaffected in C9orf72-ALS iNeurons. **(A)** Representative Western blot for BNIP3L and tubulin. **(B)** Quantification of BNIP3L expression normalised to tubulin. **(C)** Representative Western blot for BNIP3 and tubulin. **(D)** Quantification of BNIP3L expression normalised to tubulin. **(E)** Representative Western blot for C9orf72 and tubulin. * represents a non-specific band. **(F)** Quantification of NIX expression normalised to GAPDH. **(G-H)** Correlations between mitochondrial co-localisation and protein expression levels for BNIP3L (G) and BNIP3 (H) (Pearson’s correlation coefficient). Each data point represents one unique differentiation of a control/C9orf72-ALS line. All quantification was performed on 3 different differentiations of 3 control and C9orf72-ALS iNeuron lines. Data from each patient/control pair are uniquely colour-coded or labelled.

**Supplementary Figure 5:**
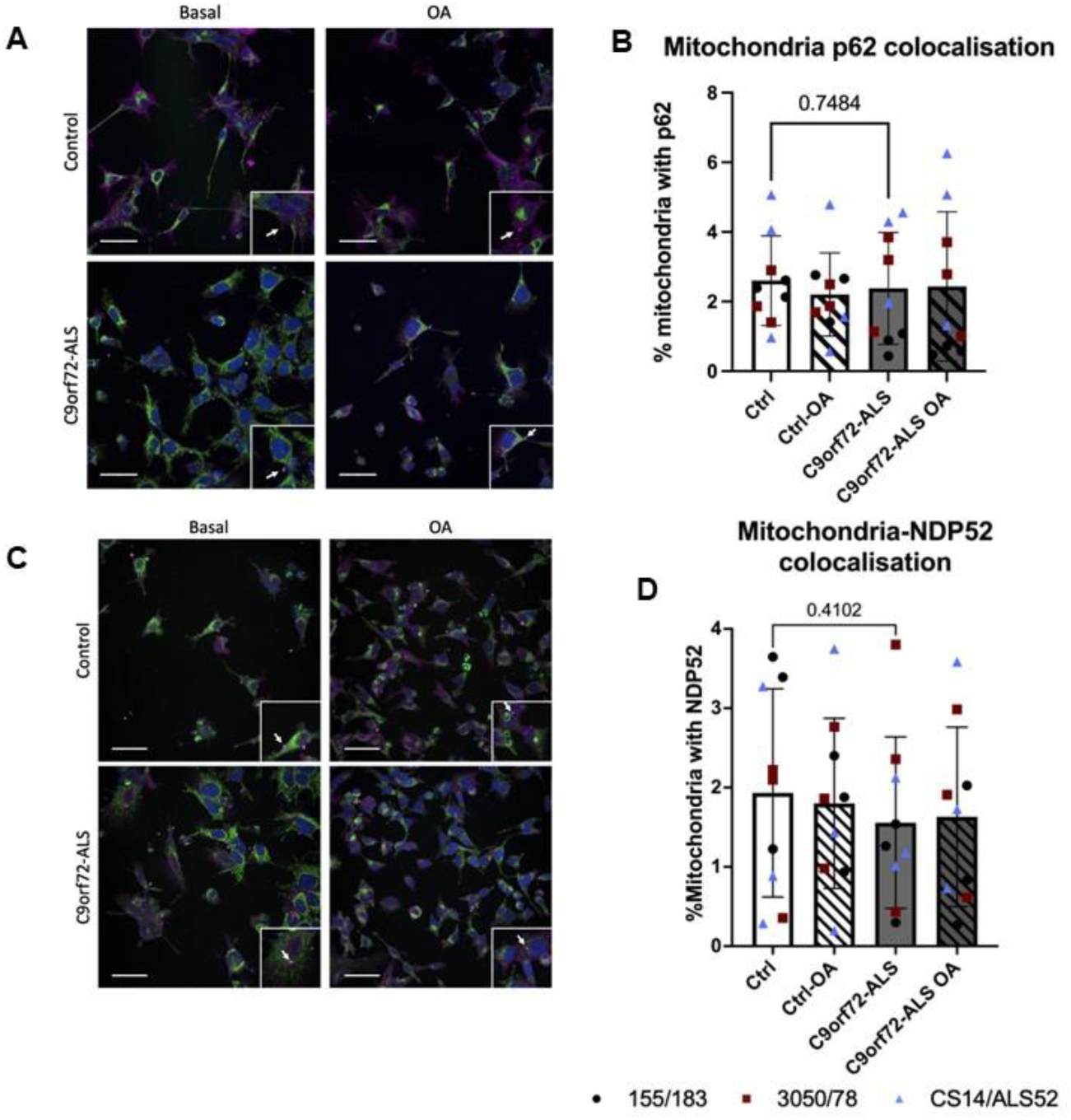
P62 and NDP52 co-localisation with mitochondria are unaffected in C9orf72-ALS iNeurons. **(A)** Representative images of control and C9orf72-ALS iNeurons stained with mitochondrial marker TOM20 (green) and p62 (magenta). Scale bar= 100μM. **(B)** Quantification of percentage of mitochondria staining with p62, under basal conditions and after mitophagy induction with oligomycin/antimycin A (OA) (mean ±SD, unpaired t-test). **(C)** Representative images of control and C9orf72-ALS iNeurons stained with mitochondrial marker TOM20 (green) and NDP52 (magenta). Scale bar= 100μM. **(D)** Quantification of percentage of mitochondria staining with NDP52, under basal conditions and after mitophagy induction with oligomycin/antimycin A (OA) (mean ±SD, unpaired t-test). Each data point represents one unique differentiation of a control/C9orf72-ALS line, taken from a mean of approximately 100-500 cells. All quantification was performed on 3 different differentiations of 3 control and C9orf72-ALS iNeuron lines. Data from each patient/control pair are uniquely colour-coded.

**Supplementary Figure 6:**
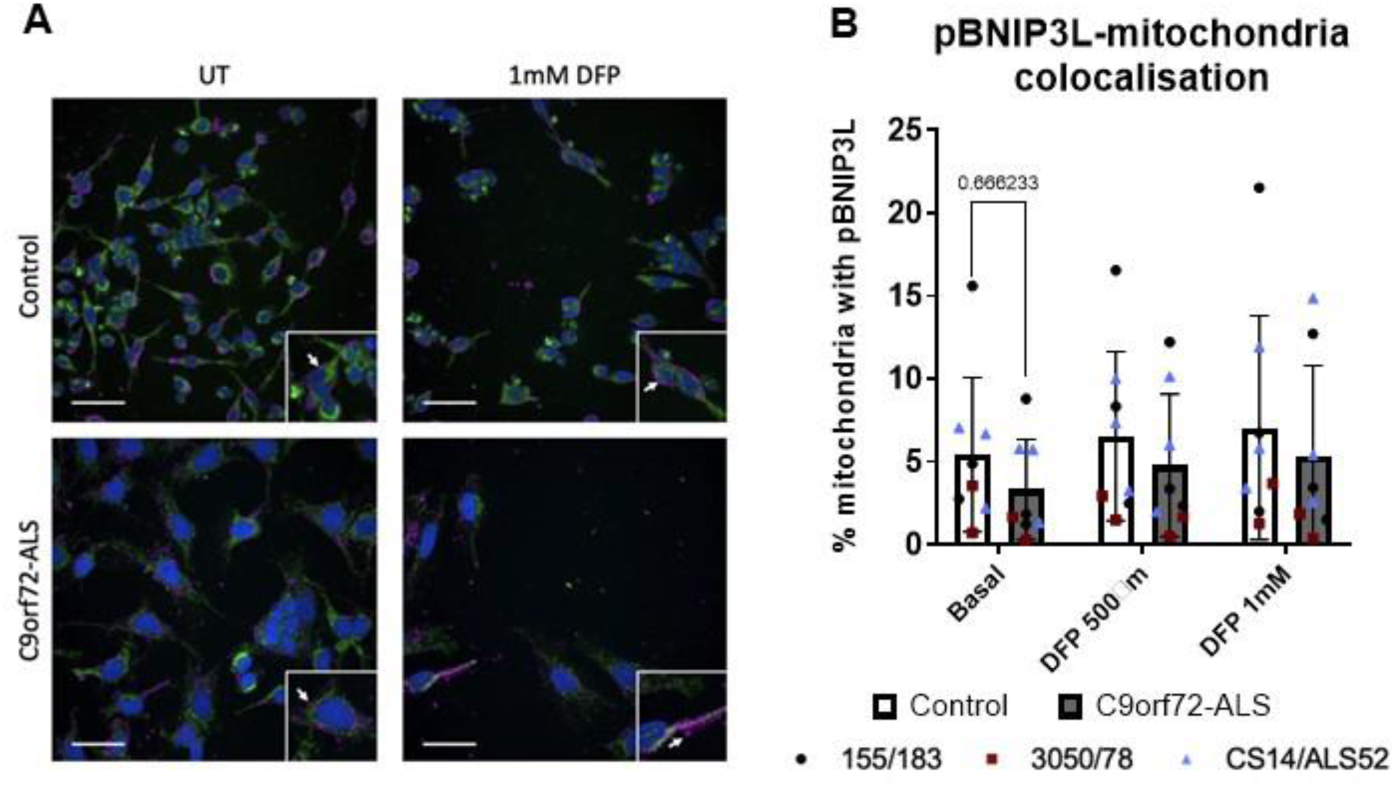
Co-localisation of phosphorylated-BNIP3L with mitochondria in control and C9orf72-ALS iNeurons. **(A)** Representative images of control and C9orf72-ALS iNeurons stained with mitochondrial marker TOM20 (green) and phospho-BNIP3L (magenta). Scale bar= 100μM. **(B)** Quantification of percentage of mitochondria staining with pNIX, under basal conditions and after mitophagy induction with Deferiprone (DFP) (mean ±SD, unpaired t-test). Each data point represents one unique differentiation of a control/C9orf72-ALS line, taken from a mean of approximately 100-500 cells. All quantification was performed on 3 different differentiations of 3 control and C9orf72-ALS iNeuron lines. Data from each patient/control pair are uniquely colour-coded.

**Supplementary Figure 7:**
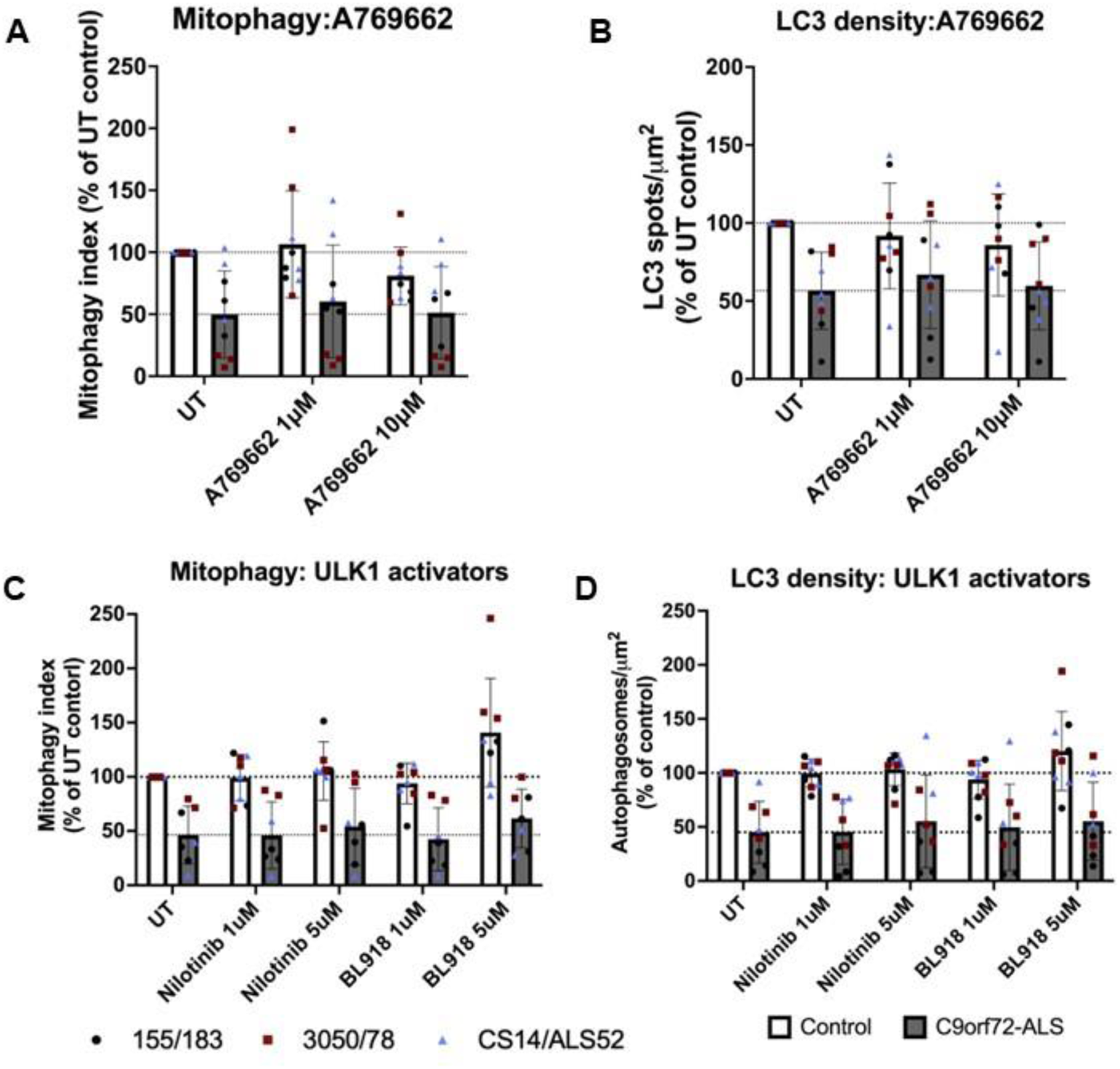
A769662, Nilotinib and BL918 do not rescue mitophagy or autophagosome production in C9orf72-ALS iNeurons. **(A-B)** Quantification of mitophagy (A) and autophagosome density (B) after A769662 treatment in control and C9orf72-ALS iNeurons (mean ±SD). **(C-D)** Quantification of mitophagy (A) and autophagosome density (B) after nilotinib and BL-918 treatment in control and C9orf72-ALS iNeurons (mean ±SD). Each data point represents one unique differentiation of a control/C9orf72-ALS line, taken from a mean of approximately 100-500 cells. All quantification was performed on 3 different differentiations of 3 control and C9orf72-ALS iNeuron lines. Data from each patient/control pair are uniquely colour-coded.

## Author contributions

JAKL: Methodology, Formal analysis, Investigation, Writing-Original Draft, Writing-Review and Editing

SG: Methodology, Investigation

KR: Methodology, Investigation

SPA: Resources, Supervision, Funding acquisition, Writing-Review and Editing LF: Resources, Writing-Review and Editing

PJS: Resources, Supervision, Funding acquisition, Writing-Review and Editing

HM: Conceptualisation, Formal analysis, Supervision, Funding acquisition, Writing-Review and Editing

